# *In silico* PK predictions in Drug Discovery: Benchmarking of Strategies to Integrate Machine Learning with Empiric and Mechanistic PK modelling

**DOI:** 10.1101/2024.07.30.605777

**Authors:** Moritz Walter, Ghaith Aljayyoussi, Bettina Gerner, Hermann Rapp, Christofer S. Tautermann, Pavel Balazki, Miha Skalic, Jens M. Borghardt, Lina Humbeck

## Abstract

A successful drug needs to combine several properties including high potency and good pharmacokinetic (PK) properties to sustain efficacious plasma concentration over time. To estimate required doses for preclinical animal efficacy models or for the clinics, *in vivo* PK studies need to be conducted. While the prediction of ADME properties of compounds using Machine Learning (ML) models based on chemical structures is well established in drug discovery, the prediction of complete plasma concentration-time profiles has only recently gained attention. In this study, we systematically compare various approaches that integrate ML models with mechanistic PK models to predict PK profiles in rats after i.v. administration prior to synthesis. More specifically, we compare a standard noncompartmental analysis (NCA) based approach (prediction of CL and V_ss_), a pure ML approach (non-mechanistic PK description), a compartmental modeling approach, and a physiologically based pharmacokinetic (PBPK) approach. Our study based on internal preclinical data shows that the latter three approaches yield PK profile predictions of comparable accuracy (evaluated as geometric mean fold errors for each profile) across a large test set (>1000 small molecules). In summary, we demonstrate the improved ability to prioritize drug candidates with desirable PK properties prior to synthesis with ML predictions.

## INTRODUCTION

Drug discovery is a multiparameter optimization problem. Therefore, ranking/prioritizing compounds based on multiple parameters is a non-trivial task. Several scores to prioritize compounds, summarizing a variety of underlying properties have been developed, e.g., ligand efficiency^1^, lipophilic ligand efficiency^2^ and Quantitative Estimate of Drug likeness (QED) Score^3^. These scores are relatively easy to calculate but suffer from insufficient transferability to the later stages of drug discovery due to missing relevance for crucial parameters, like the required dose to achieve efficacy. Predicting an efficacious dose is more challenging as it consists of two parts, namely the concentration of a drug at the target site to achieve relevant target engagement or even efficacy and the pharmacokinetics (PK), which determine the dose to achieve this required exposure. Combining both aspects in a single score was recently introduced in the Compound Quality Scores (CQS)^4^, which combine multiple key PK parameters such as the volume of distribution (V_SS_) and the clearance (CL) with the compound-specific potency readout in a single score. In this work, we focus on strengthening the PK component of these CQS, however the methodology can equally be applied to all other *a priori* PK predictions.

While individual animal or human PK parameters, such as CL or V_SS_ for molecules have been predicted by machine learning (ML) models in numerous studies^5–13^, the prediction of complete PK profiles from chemical structures (sometimes complemented by predicted or measured *in vitro* ADME data) has only been reported in a few recent studies. Handa et al trained random forest models to predict plasma concentrations in mice after i.v. and p.o. dosing for predetermined time points (one model per time point).^14^ In a different study, plasma concentrations for rats at different time points were predicted by a single model for i.v. and p.o application routes.^13^ In the aforementioned studies, PK profiles were predicted without utilizing any traditionally applied PK modeling approaches. In contrast, in other studies the input parameters to mechanistic PK models were predicted before obtaining PK profiles by conducting the PK model simulations. The PK models in those studies ranged from simple compartmental PK models (one-, two-, or three-compartment models)^15,16^ to relatively complex physiologically based pharmacokinetic (PBPK) models.^16–20^ In a modeling approach called DeepCt, rat PK was predicted using a deep learning framework that incorporates compartmental PK models.^15^ In a different study, both one-compartmental models and PBPK models were investigated to predict PK in rats after i.v. dosing.^16^ They report that AUCs of predicted profiles were of comparable quality for both approaches, although one-compartment models due to their simplicity were incapable of describing distribution of compounds to peripheral tissues. PBPK models predict the exposure in various organs based on physiological knowledge, yet their complexity limits their applicability in high-throughput scenarios. To mitigate this limitation, it has been proposed to replace PBPK models by a surrogate neural network trained to map from the inputs to the output of a PBPK model.^21,22^

The success of the reported methods has been evaluated in different ways. For example, by either directly comparing predicted concentrations with experimental concentrations or evaluating different parameters extracted from the predicted profiles (e.g., CL, V_ss_, AUC, F, c_max_, t_max_). Overall, it is challenging to compare the quality of different methodologies reported due to different predicted species (human PK or preclinical species), different training data sets, different information used for prediction (e.g., only chemical structure available vs. experimental ADME data available), different strategies for splitting training and test data (a random split typically overestimates model performance for a prospective setting^23,24^), and finally, different exposure metrics selected to evaluate the predictions (see above).

In the present study, we directly compared four different strategies for rat i.v. PK profile prediction: prediction of CL and V_ss_ by ML followed by the generation of one-compartmental models (“Baseline-ML”), direct prediction of plasma concentration by ML (“Pure-ML”), prediction of the input parameters to one-/two-compartmental models by ML (“Compartmental-ML”), prediction of the input parameters to PBPK models by ML (“PBPK-ML”). In contrast to previous studies, we evaluated the quality of prediction for the profiles with an approach that considers the entire range of the profile without being biased by overweighing earlier time points where the sampling frequency typically is larger (see Methodology). Additionally, the analysis provides a direct comparison of all methods based on a large internal and curated dataset of PK profiles for around 8k compounds.

## METHODS

### Data

For this study, we assembled in-house datasets at Boehringer Ingelheim containing *in vivo* rat PK studies with complete plasma concentration-time profiles of around 8k compounds investigated either in cocktail or single compound PK studies after i.v. application in rats (*Rattus norvegicus*). A summary of the overall compound characteristics and quality criteria for inclusion can be found in the SI.

### Study design

We tested four different PK modeling approaches and combined these with ML approaches to predict plasma concentration-time profiles in rat after i.v. administration. ML was applied to either directly predict the full profile (“Pure-ML approach”) or to predict the required input parameters for the respective PK model. All four PK modeling approaches differ in the degree of mechanistic representation of the distribution, metabolism, and excretion processes. Pure-ML is a purely empiric approach and is applied directly to predict plasma concentrations without any interpretation of the underlying PK processes. The “Baseline-ML” includes key PK parameters such as V_SS_ or CL but does not include a more detailed description of the shape of the PK profile, i.e., a standard one-compartment PK model is applied. The next level of mechanistic representation is an integrated one-/two-compartment PK approach, which can additionally account for distribution between the central and peripheral compartment and is a commonly applied approach in PKPD modelling. Finally, the PBPK-ML approach relies on a more mechanistic representation of the organism by considering physiological parameters such as blood flows, organ volumes, and tissue compositions to predict the distribution into tissues based on physico-chemical properties of a compound.^25^

We employed a temporal splitting strategy^23^ to critically evaluate the predictive performance of all four approaches in a real-life application by drug discovery project teams. We trained multiple models for each approach based on data that would have been available at a certain point in time. For instance, the first model was trained on data for all compounds registered before the end of the year 2017 and used to make predictions for compounds registered in the first three months of 2018. Next, the data for the test set compounds of the first model (January to March 2018) was added to the training set for the second model and this model again was used to predict PK profiles for compounds in the following three months (April to June 2018). In this way 18 models were trained until predictions for the most recent compounds of the dataset were made. This principle is illustrated in Figure 1. For all summary statistics and general evaluation of the prediction quality, all PK predictions for all 18 test sets were pooled (pooled test set).

**Figure 1.**
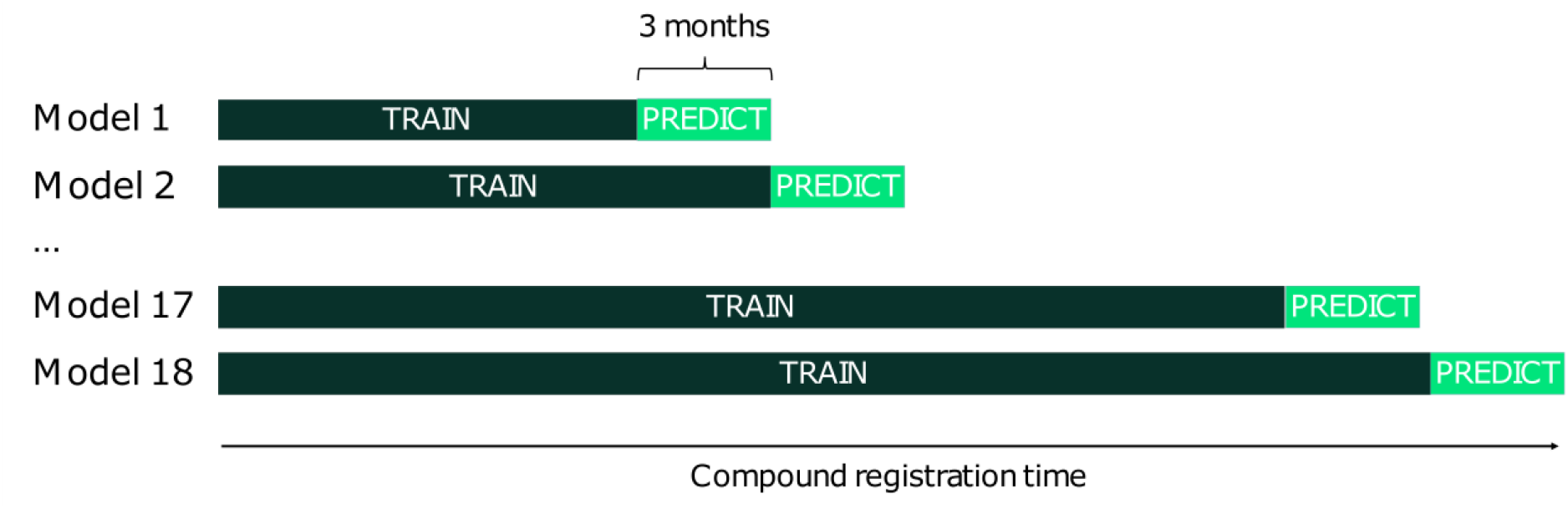
Temporal splitting strategy.

### Baseline-ML

The baseline to our current efforts was inspired by a recently published study on CQS.^4^ This study was geared towards providing a single score allowing compound prioritization by combining PK properties and *in vitro* compound potency. In this approach the PK has been described by a one-compartment model with instantaneous absorption of the required dose. In the first step, NCA parameters (CL and V_SS_) were estimated. Then, two separate ML models were trained to predict the NCA parameters (i.e., CL and V_ss_). Finally, the predicted parameters were used to generate PK profiles.

The ML model used as input 4414 molecular descriptors (selected AlvaDesc descriptors)^12,26^ and 10 predicted ADME descriptors. Those ADME descriptors were predicted by a different ML model beforehand (for more details see SI). Following the implementation in the CQS publication, the extremely randomized trees algorithm with 600 estimators was used as ML model. The approach is summarized in Figure 2 (left).

**Figure 2.**
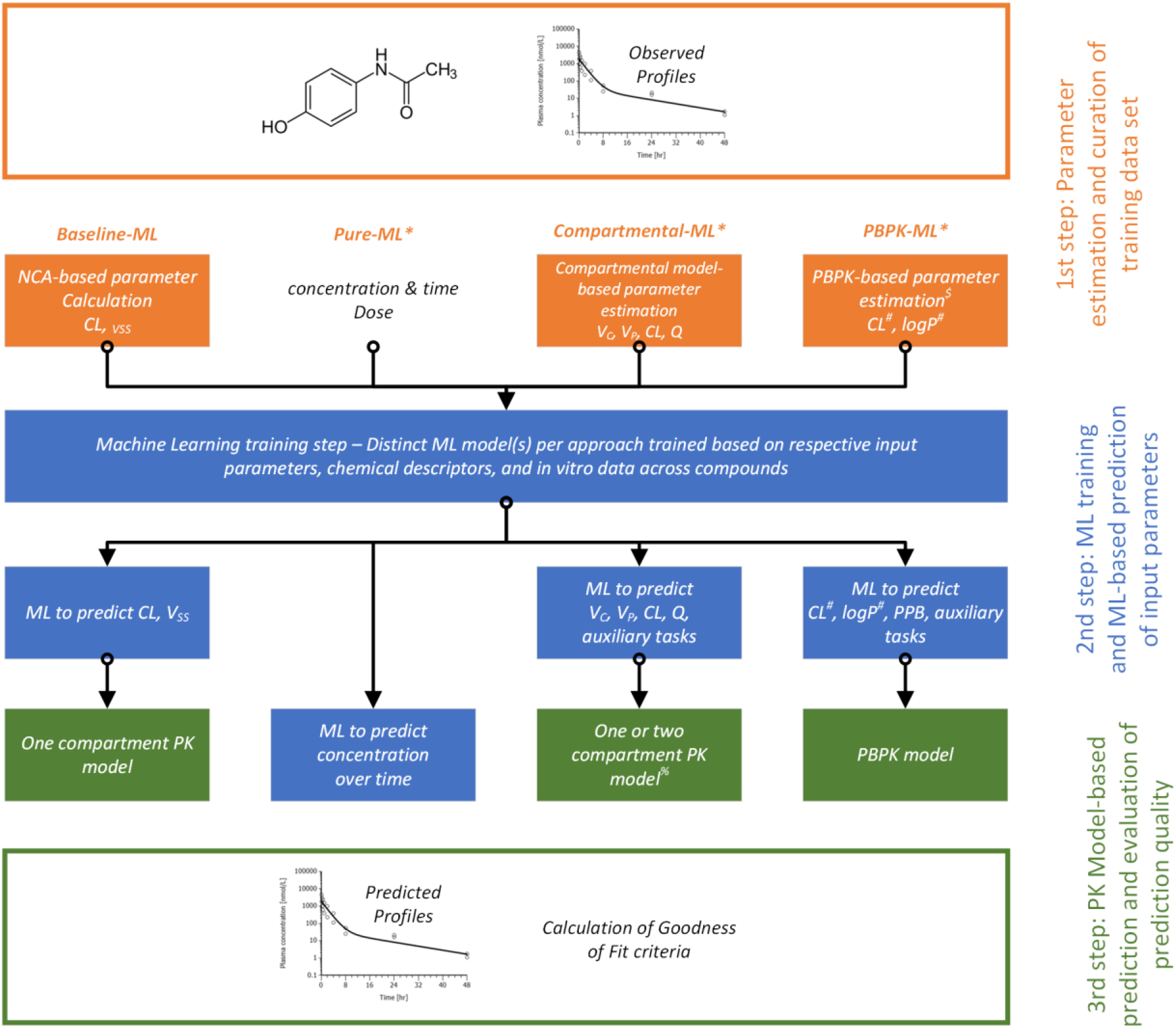
Overview of PK profile prediction methods (details in SI). *Pure-ML, Compartmental-ML, and PBPK-ML were new methodologies, which were compared to the Baseline-ML, which was previously part of the Compound Quality Scores. #CL, and logP values are input parameters for the PBPK model and should not be mixed with predicted parameters for the compartmental model or respective in vitro data (i.e., surrogate parameters for elimination and distribution in the PBPK model). $Input parameters such as PPB to the first step were either predicted or based on existing in vitro data (e.g., the PPB, CL, logP) as input or also as initials for the parameter estimation.

### Pure-ML

The method establishes a benchmark for our ability to forecast compound plasma concentration without applying any compartmental models (see Figure 2, middle-left). The Pure-ML model was trained to forecast the logarithm of compound plasma concentration at 6 minutes intervals. With the trained Pure-ML model, plasma concentration can be predicted by providing a representation of the molecule of interest (see below), a dose and the time point of interest.

In the Pure-ML model, each molecule was represented as a set of molecular descriptors (4414 selected AlvaDesc descriptors^12,26^) and 12 predicted ADME/PK features. Those ADME/PK features were predicted by a separate ML model beforehand (see SI). In addition, dose and the time point were used as features in the Pure-ML model. As ML algorithm, a LGBM regressor^27^ (version 3.2.1) was used with default parameters except for setting number of estimators to 1000.

### Compartmental-ML

To estimate the true “optimal” PK parameters for all compounds in the dataset, an automated fitting algorithm was developed. In a first step, initial parameters were identified by noncompartmental analysis. Parameter estimation was then performed in a second step using the nlmixr package within R utilizing the initial estimates for k_el_, k_32_, k_23_, V_c_ and V_p_ (with k_23_ and k_32_ fixed to zero in the case of a one compartment model fit). These parameters can also be directly transformed into the more common parameters, namely CL, central volume of distribution (V_c_), peripheral volume of distribution (V_p_), and the intercompartmental CL (Q). For compounds which were adequately described with a one compartment PK model, V_p_ was set to a very small value (0.01 L/kg), and Q was set to a high value (12 L/h/kg) (which is basically identical to a one-compartment PK model). This ensures that no model selection is necessary in the ML step. A logarithmic transformation (with the base 10) was applied to all the compartmental parameters before training ML models. A single ML model was trained on the four compartmental parameters (multi-task architecture, see below). Finally, a molecule can be provided to the Compartmental-ML model to predict the four parameters from which a PK profile can be calculated.

The multi-task ML model was trained to predict the compartmental parameters as main (i.e., target) tasks, as well as to predict auxiliary *in vitro* ADME and *in vivo* PK tasks. This was motivated by a previous study revealing that prediction accuracy may be improved when target tasks are co-learned with auxiliary tasks that are at least weakly related to the target tasks.^12^ The ML model was trained using the Chemprop package (version 1.5.2) in Python, which implements neural networks that may learn directly from chemical graphs as input using the message-passing framework.^28^ The approach is summarized in Figure 2 (middle-right). A more detailed description of the method is provided in the online supplementary material.

### PBPK-ML

In the PBPK-ML approach (see Figure 2, right), a generic PBPK model for small molecules was developed in PK-Sim^29^ as part of the Open Systems Pharmacology Suite, Version 11.1.^30,31^. A mean rat individual weighing 227 g was generated with the PK-Sim physiology database and used for all simulations. Partition coefficients and cellular permeabilities were calculated using the Berezhkovskiy and PK-Sim standard calculation methods, respectively. Clearance of the compound was selected as “linear liver plasma CL” in PK-Sim. No renal elimination or more specific (hepatic) metabolization by specific enzymes were considered. As not all parameters were measured for all compounds *in vitro* (especially plasma protein binding), the missing parameters had to be predicted as input parameters already prior to the machine learning step.

In a second step, after parameterizing the PBPK model with using *in vitro* or *in silico* parameter values only, fitting of lipophilicity and liver plasma CL to observed plasma concentration profiles was performed. Best parameter values were estimated using the parameter identification package {ospsuite.parameteridentification}^32^ implemented in R. A combination of global and local optimization algorithms with the M3 error model was applied. These estimated “optimal” lipophilicity and “liver plasma CL” values were then combined with other available data to train a multi-task ML model comparable to Compartmental-ML based on chemical structures. Finally, PK profiles were simulated using predicted lipophilicity, liver plasma CL, PPB from the ML model, pKa values from MoKa (version 2.6.0)^33^, and other required input parameters from the chemical structures. It is important to highlight that these CL and lipophilicity values do not represent predicted values of *in vitro* assays, but of input parameters previously estimated as “optimal parameters”.

In addition to the required input for PBPK simulations (lipophilicity, liver plasma CL, PPB), the PBPK-ML model was trained on auxiliary tasks in a comparable manner as for Compartmental-ML (for details see SI).

### Evaluation of predicted PK profiles

To evaluate the quality of predicted profiles, quite often the predicted plasma concentration-time profile is compared with the observed plasma concentrations. However, sampling in PK studies is typically more frequent at early time points compared to later time points. Hence, evaluating the predictive performance solely based on observed data would bias the evaluation strongly towards early time points. Instead, we evaluated the predicted concentration-time profiles of all four approaches against fitted profiles (one-/two-compartment fits), which allows an interpolation between all observed data points. In particular, the geometric mean fold error (GMFE) was calculated for each predicted profile. The GMFE is defined as:

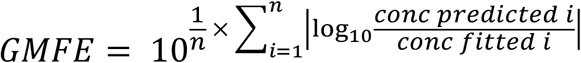

where predicted and fitted plasma concentrations were considered in intervals of 0.1 h (i.e., 6 min) up to the last time point where an experimental measurement exceeded the lower limit of quantification (LLOQ). The principle is illustrated in Figure 3, although for simplicity in the plot fold errors are only shown at full hour time points. As the last experimental time point for this compound was obtained at t=8h, the calculation of the GMFE stops at this point. At the latest, calculations were stopped at t=24h.

**Figure 3.**
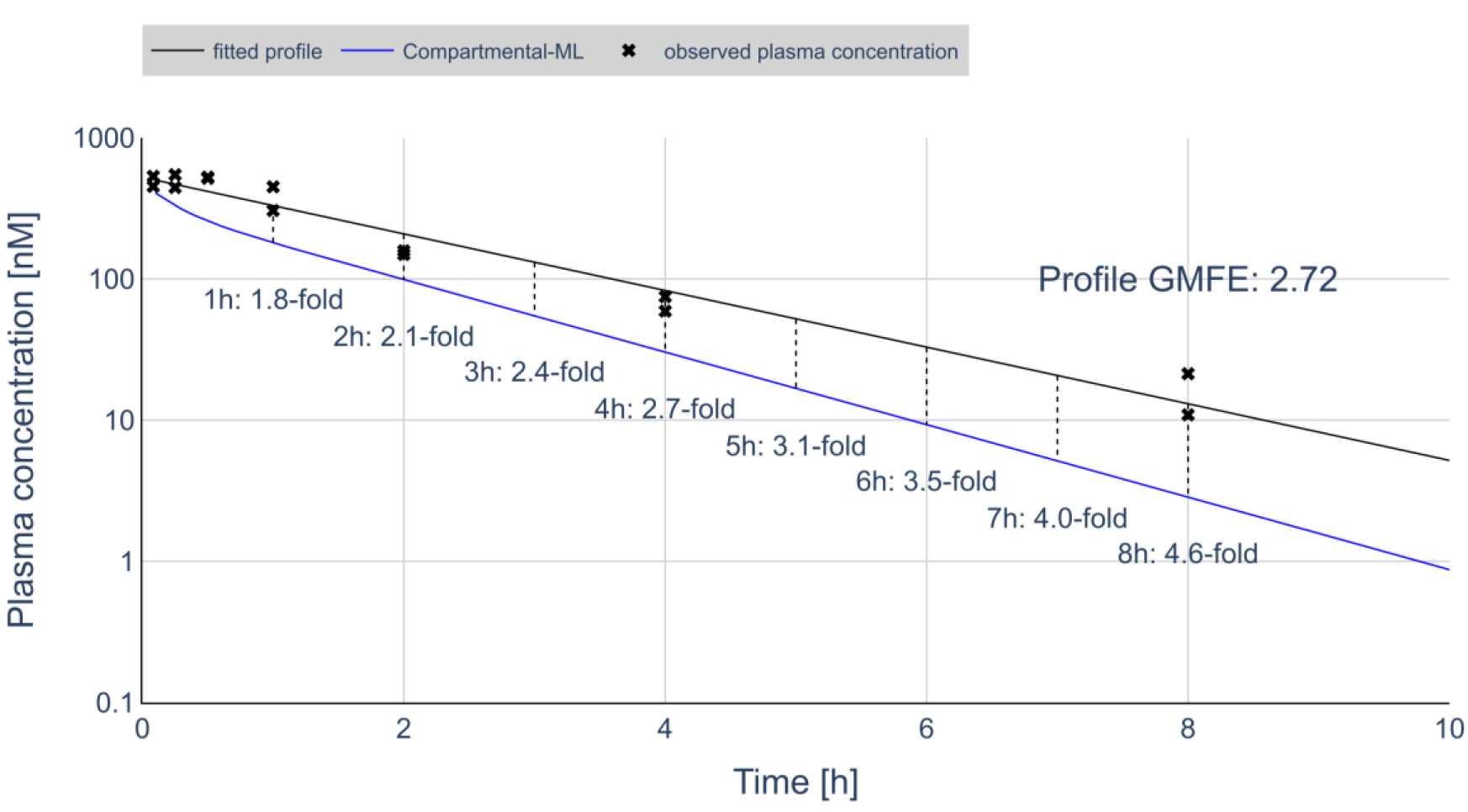
To evaluate the quality of predicted PK profiles, a GMFE is calculated by considering concentrations in time intervals of 6 minutes. Shown in the plot are the fold errors at t= [1h, 2h, …, 8h] (1h intervals for illustration) as well as the overall GMFE calculated using the 6 min intervals.

We further analyzed the magnitude and bias of fold errors for the different approaches over time with a visualization related to visual predictive check (VPC), which is a popular tool for population PK studies.^34^ For that purpose, we calculated the 5^th^,10^th^, 25^th^, 50^th^, 75^th^, 90^th^ and 95^th^ percentiles of the ratio of predicted to fitted plasma concentrations at each time point (every 6 minutes from 0 to 8 hours) across all compounds in the test set. These ratios are then overlaid as bands over a typical PK profile generated as the median of all the fitted PK parameters (including training set). The median of the ratio between the true PK simulation and predicted simulation is shown as a solid line while the different percentiles are shown as bands that display the overall deviation of prediction from truth over time.

## RESULTS

### Performance of PK profile prediction

We used the GMFE as a metric to evaluate the quality of predicted profiles. The summary of obtained GMFE values for the pooled test set of 1217 compounds for all four prediction approaches is provided visually as a cumulative distribution plot in Figure 4. Three of the methods (Compartmental-ML, PBPK-ML, Pure-ML) achieved comparable median GMFE values slightly below 3-fold and no method seems to be clearly superior to the others due to the highly similar distributions of GMFEs, whereas Baseline-ML overall performed much worse with a median GMFE of 4.49-fold (see Table S3). This approach was therefore clearly incapable to accurately predict the profiles. The Baseline-ML approach assumes a monophasic decline of the plasma concentration-time profile, whereas all other approaches allow for multiphasic distribution and elimination. That indicates that this modelling feature is important to achieve a higher prediction accuracy. Among those three approaches, Pure-ML achieved the lowest number of predictions with very high GMFE values (e.g., only 1.3% of predictions with GMFE>100, 3.4% for PBPK-ML, 4.9% for Compartmental-ML).

**Figure 4.**
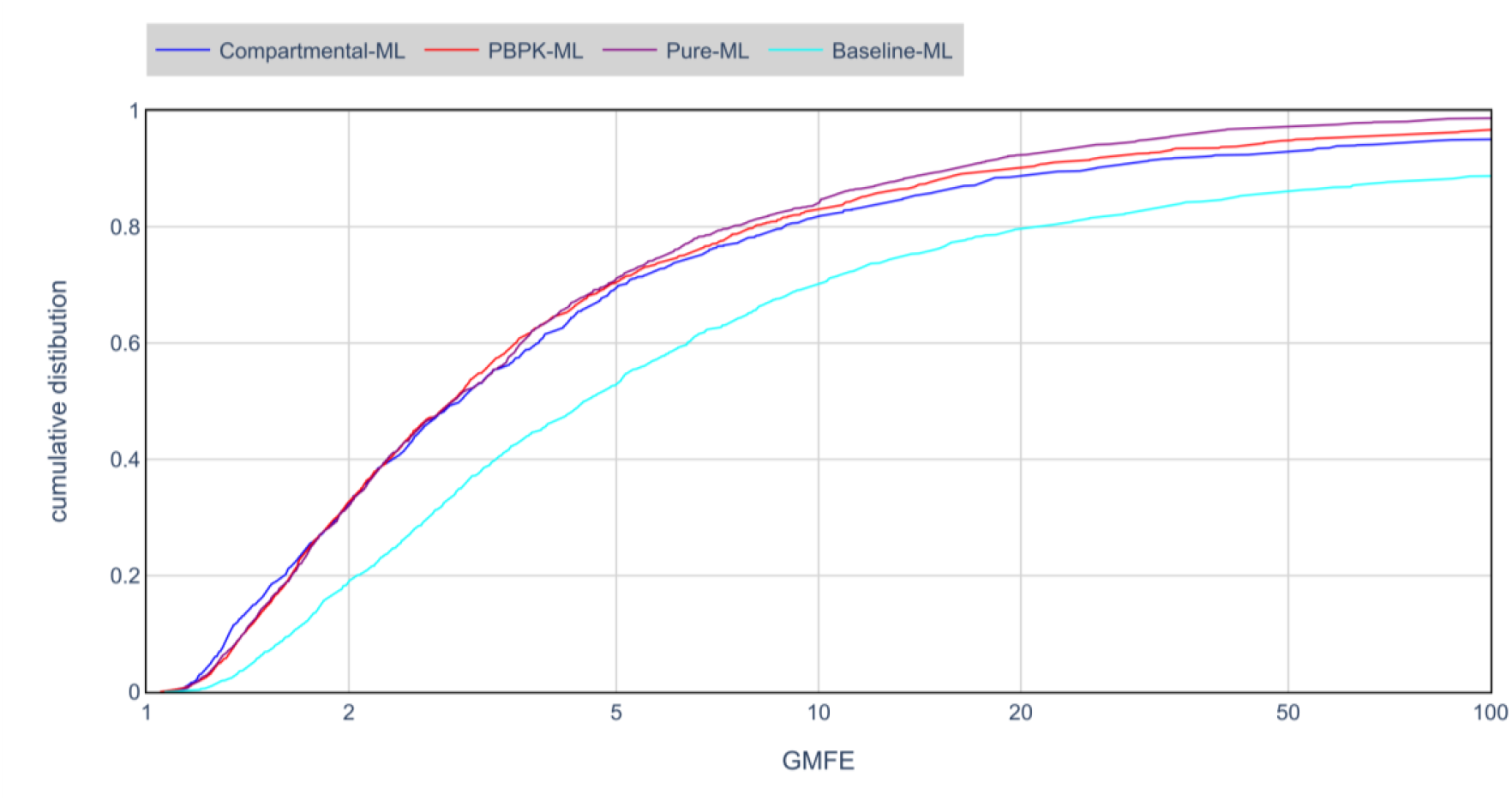
Cumulative distribution of GMFE values for each method.

A few representative examples of predicted versus fitted profiles are shown in Figure 5. The first row (Examples 1 to 3) shows compounds with rather short terminal half-lives (last experimental measurement after 2 or 4 hours), the second row (Examples 4 to 6) comprises compounds with measurements above the lower limit of quantification (LLOQ) for 8 hours after dosing, and the third row (Examples 7 to 9) contains compounds with longer half-lives, which means that the last observation after 24 hours or later is still above the LLOQ. Within each row, examples were selected to be highly accurate predictions (first column), predictions with moderate accuracy (second column), as well as poor predictions (third column). The GMFEs for all predicted profiles are given in the caption of Figure 5. Note that Baseline-ML always generates a linear profile (on a logarithmic plasma concentration scale), whereas the three other approaches can predict more flexible profiles that can more accurately reflect fitted PK profiles. Profiles predicted by the Pure-ML approach do not decrease perfectly monotonically. This is because each time point is predicted independently by the ML model, and hence no mechanistic PK parameterization leads to a continuous decrease of plasma concentration over time. Nonetheless, the overall shapes of predicted profiles resemble those of typical fitted PK profiles. As additional information, we visualize the correlation of GMFE scores between each pair of methods (Figure S1). Briefly, some correlation exists between the GMFEs achieved by different methods for a particular compound suggesting that based on our data some compounds are more challenging to predict than others. However, examples exist where the quality of prediction differs strongly between two methods.

**Figure 5.**
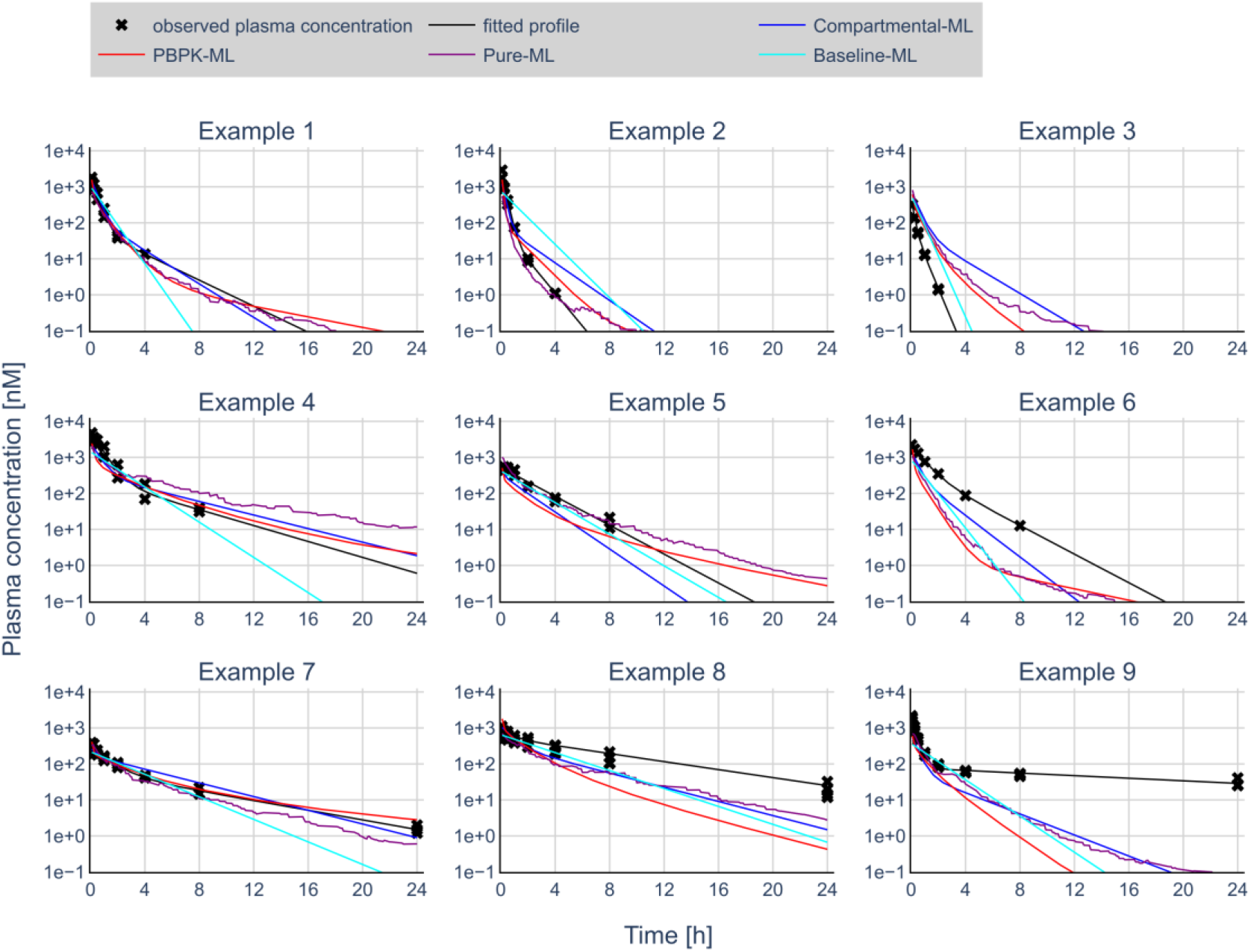
Nine representative examples illustrating predicted versus observed plasma concentration-time profiles. The first row shows compounds with rapid elimination (i.e., last observed time point at t=2h or t=4h), the second-row compounds with last observed time point at t=8h, and the third row compounds with longer half-lives, i.e. last observed time point is at t=24h. Within each row, the examples are sorted according to quality of predictions from lower to higher GMFEs (given in brackets below). **Example 1**: Compartmental-ML (1.30), PBPK-ML (1.28), Pure-ML (1.24), Baseline-ML (1.54); **Example 2**: Compartmental-ML (2.69), PBPK-ML (2.13), Pure-ML (2.16), Baseline-ML (9.28); **Example 3**: Compartmental-ML (8.65), PBPK-ML (5.29), Pure-ML (6.95), Baseline-ML (5.54); **Example 4**: Compartmental-ML (1.50), PBPK-ML (1.50), Pure-ML (2.12), Baseline-ML (1.43); **Example 5**: Compartmental-ML (2.72), PBPK-ML 2.86), Pure-ML 1.31), Baseline-ML (1.50); **Example 6**: Compartmental-ML (3.91), PBPK-ML (18.1), Pure-ML (12.2), Baseline-ML (10.3); **Example 7**: Compartmental-ML (1.36), PBPK-ML (1.23), Pure-ML (1.78), Baseline-ML (4.11); **Example 8**: Compartmental-ML (5.01), PBPK-ML (12.3), Pure-ML (4.50), Baseline-ML (5.76); **Example 9**: Compartmental-ML (45.6), PBPK-ML (230), Pure-ML (37.6), Baseline-ML (159).

We further analyzed trends in prediction error over time for the different prediction approaches. Figure 6 shows a VPC comparison across predicted and observed concentration-time profiles (i.e., fitted to the raw data) in the time span of 0h to 8h (0h to 24h in Figure S2). It illustrates the accuracy and bias of the predictions in relation to a “normalized” PK profile for our dataset (details in Methods and caption of Figure 6). For Compartmental-ML and Pure-ML, the median of all predictions across the test set is nearly identical to the observed median across the entire time span (here 0 to 8 hours), indicating no bias (i.e., no systematic over- or underprediction). For all methods, the accuracy of predictions for initial time points falls into a narrow range, whereas for later time points progressively more extreme over- and underpredictions occur. For instance, for Compartmental-ML the 25^th^ and 75^th^ percentiles at t=0.5h indicate a 1.4-fold under- and a 1.9-fold overprediction, whereas the numbers increase to a 4.6-fold under- and 6.0-fold overprediction at t=8h. PBPK-ML shows a bias towards underprediction at later timepoints with a median fold error of 1.9-fold at t=8h. Baseline-ML shows a bias of overpredicting early time points (1-3 hours) and a systematic underprediction of 4.5-fold at t=8h. This is in line with the application of a one-compartment PK model, which does not account for steeper initial distribution and flatter terminal elimination phases. Overall, this analysis revealed that the lowest biases occurred for Compartmental-ML and Pure-ML. For all the methods, predicted concentrations deviate more strongly from fitted profiles at later time points.

**Figure 6.**
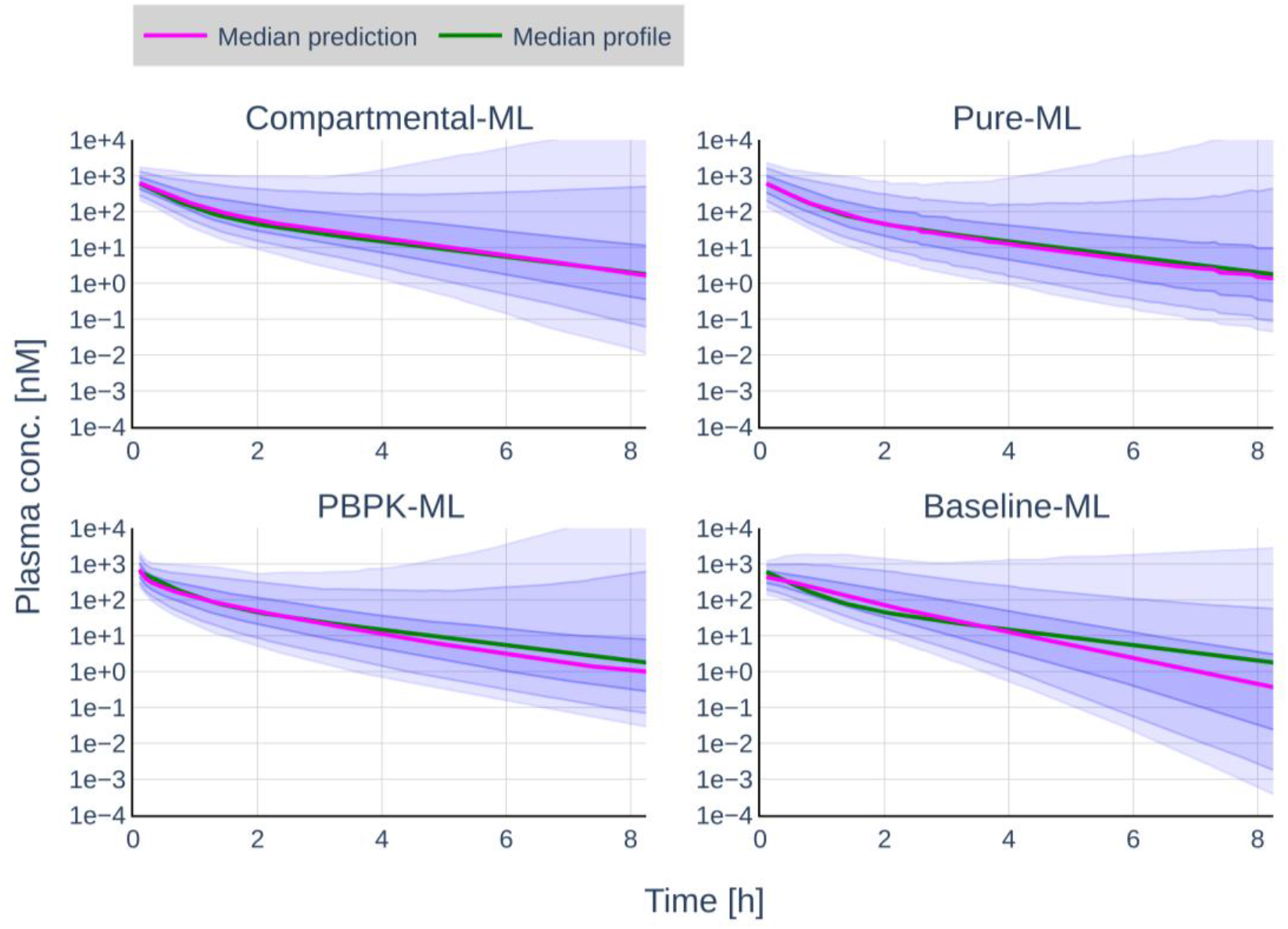
Visualization of prediction accuracy and bias over time. At each time point (0 to 8 hours in 6-minute intervals) the ratios of predicted concentrations to fitted concentrations were calculated. The plots show a typical profile generated from median parameters across the dataset (green line), the median deviation from a typical profile (50^th^ percentile, pink line), as well as (from top to bottom) the 95^th^, the 90^th^, the 75^th^, the 25^th^, the 10^th^, the 5^th^ percentile of fold errors for each method at the different time points.

## DISCUSSION

The presented analysis herein demonstrates that full plasma concentration-time profile predictions based on different PK modelling approaches can provide additional value beyond the standard *in silico* PK predictions, which are mainly based on CL and V_SS_ prediction. Compared to Baseline-ML, the other methods achieved higher accuracy (evaluated as GMFE) as well as lower bias in their predictions. To our knowledge, this work also represents the first systematic comparison of PK modelling approaches in combination with machine learning to assess which is best suited for *a priori* predictions of plasma concentration-time profiles.

For all methods investigated here, conceptually comparable methods have been previously reported in the literature (related studies to each method: Baseline-ML^16^, Pure-ML^13,14,35^, Compartmental-ML^15^, PBPK-ML^16,18–21^), although modeled species, application routes, and details of the ML method (e.g. input features, ML algorithm) may differ. Therefore, we consider our study to cover a wide range of methodological concepts employed also by other researchers. A direct comparison of our results to the literature is not possible, as other studies did not systematically evaluate accuracy across complete profiles (see Methods). We also note that *in vivo* PK data usually are proprietary, and no suitable public benchmarks exist to compare different approaches.

All three methods, namely PBPK-ML, Compartmental-ML, and Pure-ML, yielded comparable results, all of which outperformed the Baseline-ML approach. As all three prediction methods for full concentration-time profiles provided comparable prediction quality, this might indicate that the predictive power might not be limited by the respective modelling approach, but rather by how well the ML step can capture the relationship between the chemical structure and the respective input parameters (e.g., both logP, CL for PBPK-ML or CL, V_c_, V_p_, Q for Compartmental-ML). Therefore, a more detailed discussion of the different methods might be necessary to determine the individually preferred approach.

Overall, the results indicate that the Pure-ML approach has the smallest median GMFE for predicting very late time points. Especially for predictions beyond 8 h observation time frames, the higher percentiles were less variable compared to the other methods (compare online supplementary). However, as the human PK is normally slower due to allometric relations, the relevance of these time frames in rats for accurately anticipating the human situation can be debated. The Pure-ML model is directly trained on the profile, including the later time points, which might explain this improved performance. In contrast, all other approaches are not directly trained on the profile but rather on the estimated input parameters. On the other hand, the drawback of this characteristic is that the Pure-ML approach does not include concrete parameters such as CL, V_SS_, making it more challenging for MedChem or DMPK scientists to interpret the results of this approach. Furthermore, this missing parameterization could also complicate scaling the rat PK profile to human.

The Compartmental-ML approach yielded very comparable results to the other full profile prediction approaches, i.e., outperforming the Baseline-ML approach. For Discovery Research scientists, especially for MedChem and DMPK scientists the parameterization with CL and V_SS_ is probably the most familiar. Furthermore, scaling of the respective parameters to human is feasible, and the technical implementation is most straightforward, for instance allometric scaling or hepatocyte-based correction between rat and human CL. Furthermore, one simple but potentially important advantage is that the prediction of the plasma concentration-time profile can be performed with a simple closed form equation for a two-compartment model with or without oral absorption. This might become of special importance when the predictions are provided in an automated (technical) framework, so that predictions are easily available to the project teams.

Out of the three investigated approaches, the PBPK-ML approach is the most mechanistic. Numerically, it had the lowest median GMFE, however the results were comparable across all three methods. Also, in comparison to the Compartmental-ML approach, slightly less outliers were detected based on the determined GMFE, potentially due to some mechanistic constraints, which might prevent some “unrealistic” PK parameterization (e.g., CL way beyond the hepatic blood flow). The main strength of the PBPK approach related to early PK predictions is the ability to scale between different species, individuals, or special populations. The human PK can be predicted with parameters estimated with preclinical data, allowing the calculation of the first in human dose.^36^ However, the high mechanistic complexity comes at the cost of the most complex structural model. No closed form equation, such as the Bateman function, is available for these models, so *a priori* predictions need to be done in a dedicated script. Furthermore, the initial investment to develop and qualify template models might be more complex, for instance additional explorations might be necessary how to include renal or extra-hepatic metabolism in a routine high-throughput workflow.

In conclusion, we have shown that it is possible to perform *a priori* predictions of complete plasma concentration-time profiles, using any of the three investigated PK-modeling methods combined with ML. Although all approaches are comparably predictive, we believe that the choice of PK modelling method should be determined by how well it can be integrated into the respective data and workflow structure. Especially, considering the high level of automation required for calculating and providing high throughput *in silico* PK and related compound quality score predictions in Drug Discovery. According to PK-based ranking results across drug discovery projects, we believe that *in silico* PK predictions have evolved into a tool that can be used to prioritize and thereby identify compounds with the best PK properties. The relevant exposure metric, such as specific surrogate time points or coverage over a certain time can be freely chosen based on what is most meaningful for establishing an efficacious human dose estimate. When these plasma concentration-time profile predictions are combined with meaningful potency predictions, they can serve as a powerful framework for prioritizing compounds and ultimately identifying a meaningful number of compounds for clinical candidate investigation.

The next phase of this research will involve demonstrating a sufficient predictive power after p.o. administration and examining whether the integration of data from multiple species (in the training data) can further improve the prioritization of compounds. Furthermore, the potential to link additional metrics such as an area under the effect curve (AUEC)^37^ or even a comprehensive pharmacodynamic (PD) model to these PK predictions, as exemplified by a previous work by Chen et al.^38^, needs to be explored. Finally, the impact and the convenience of applying these predicted PK profiles and the related CQS needs to be proven in a prospective evaluation by drug discovery project teams.

## Acknowledgements

We would like to thank the groups and teams at Boehringer Ingelheim who generated the data in this study, namely Research PK, *In vitro* ADME, Bioanalysis, and CMC as part of the global Drug Discovery Sciences Department. Furthermore, we would like to thank the global Medicinal Chemistry department for providing compounds, which went into this systematic analysis.

## Author contributions

Conceptualization: M.W., L.H., M.S., J.M.B., C.T. Data collection and curation: M.W., L.H., M.S., B.G., H.R., G.A. PBPK model development: B.G., P.B., Compartmental model development: G.A., Pure machine learning approach: M.S. Development and Implementation of ML models: M.W., M.S. Data analysis: M.W., G.A. Writing: M.W. J.M.B., L.H., M.S, G.A., B.G., H.R., C.T., P.B.

