## Supplemental Material for "*In silico* PK predictions in Drug Discovery: Benchmarking of Strategies to Integrate Machine Learning with Empiric and Mechanistic PK modelling"


#### Author info

**Moritz Walter[1], Ghaith Aljayyoussi[2], Bettina Gerner[2], Hermann Rapp[2], Christofer S. Tautermann[1], Pavel Balazki[3], Miha Skalic[1], Jens M. Borghardt[2#], Lina Humbeck[1#]**

**Affiliations**

**1: Boehringer Ingelheim Pharma GmbH & Co. KG, Medicinal Chemistry, Computational Chemistry, Biberach, Germany**

**2: Boehringer Ingelheim Pharma GmbH & Co. KG, Drug Discovery Sciences, Preclinical PKPD Modelling and Data & Digital Sciences, Biberach, Germany**

**3: ESQlabs GmbH, Saterland, Germany**

**# Corresponding authors: Lina Humbeck (, phone: +49 (7351) 54-175756, address: Birkendorfer Str. 65, 88397 Biberach, Germany) for all ML-related aspects of the manuscript, Jens M. Borghardt (, phone: +49 (7351) 54-141046, address: Birkendorfer Str. 65, 88397 Biberach, Germany) for PK modelling and DMPK aspects of the manuscript**


#### Details on the study data

Before performing the analysis, the in vivo data set was cleaned to remove non-relevant data. For example, any data from pretreated animals were removed (i.e., to remove preclinical drug-drug interaction studies). Furthermore, only studies with at least three measured time points (in at least one of the animals in the study) were included in the final dataset. When multiple studies for a given compound were available, priority was given to the study with the highest number of time points and then studies with the lowest dose. In the end, the *in vivo* dataset was a collection of 157593 *in vivo* PK data points across 8043 compounds in *Rattus norvegicus*. 25589 values were reported as below their respective reported lower limits of quantification.

#### Details on the PK model used in Compartmental-ML

For each profile an automated NCA analysis is performed to estimate the initial parameters. Using linear regression, four initial parameters are calculated: A, $\alpha$ are the intercept and slope of the first two time-points of the i.v. profile calculated on a log scale and B, $\beta$ are the intercept and slope of the last three time-points of the i.v. profile. Based on these values the constants $k_{32}$, $k_{el}$ and $k_{23}$ are calculated according to equations 1-3 for initial estimates of a two-compartment model. Vc is calculated according to equation 4.

For the initial estimates of a 1-compartment model, an additional regression is performed over the log-transformed data generating the parameters L and λ, representing the intercept and slope. Area under the curve (AUC) is calculated using a standard linear up log down method using the trapezoidal method, and its initial and terminal extrapolations ($AUC_{0\text{-}C1}$, $AUC_{Clast - inf}$) are calculated according to equations 5 and 6. Initial estimate for $V_c$ is calculated according to equation 7 while $k_{23}$ and $k_{32}$ are fixed to 0. Data fitting is then performed utilizing the nlmixr package within R using the initial estimates for $k_{el}$, $k_{32}$, $k_{23}$ and $V_c$ as previously described (with $k_{23}$ and $k_{32}$ fixed to zero in the case of a one compartment model fit). The fit with the lower value of Akaike is then chosen to represent the true pharmacokinetic parameters for that compound. Secondary parameters, CL, Q and $V_p$ are calculated according to equations 11-13.

$$k_{32} = \frac{A \cdot \beta + B \cdot \alpha}{A + B} \qquad \text{Eq... 1}$$

$$k_{el} = \frac{\alpha \cdot \beta}{k_{32}} \qquad \text{Eq... 2}$$

$$k_{23} = \alpha + \beta - k_{32} - k_{el} \qquad \text{Eq... 3}$$

$$V_c = \frac{Dose}{A + B}, two\ compartment \qquad \text{Eq... 4}$$

$$AUC_{0-Cfirst} = \frac{T_{first}}{\ln\left(\frac{A}{C_{first}}\right)} \cdot (A - C_{first}) \qquad \text{Eq... 5}$$

$$AUC_{Clast-\infty} = \frac{C_{last}}{k_{el}} \qquad \text{Eq... 6}$$

$$V_c = \frac{Dose}{AUC \cdot \gamma}, one\ compartment \qquad \text{Eq... 7}$$

$$\frac{d_{plasma}}{dt} = -(k_{el} + k_{23}) \cdot plasma + k_{32} \cdot periph \qquad \text{Eq... 8}$$

$$\frac{d_{periph}}{dt} = k_{23} \cdot plasma - k_{32} \cdot periph \qquad \text{Eq... 9}$$

$$Conc = \frac{plasma}{V_c} \qquad \text{Eq... 10}$$

$$CL = V_c \cdot k_{el} \qquad \text{Eq... 11}$$

$$Q = k_{23} \cdot V_c \qquad \textit{Eq... 12}$$

$$V_p = \frac{Q}{k_{32}} \qquad \textit{Eq... 13}$$

**Details on the PK model used in PBPK-ML**

In the PBPK-ML approach, a generic small molecule compound was utilized to establish a generic PBPK rat model. This model was developed using PK-Sim[1] as part of the Open Systems Pharmacology Suite (OSPS) Version 11.1.[2,3] The software offers a comprehensive PBPK model structure, including physiological and anatomical parameters of the relevant species, and ordinary differential equations representing the most relevant organs for the ADME process. The final ordinary differential equations model can be exported to R and solved using the ospsuite-r[4] R package.

A framework has been developed in R that allowed automated parametrization of the template model for selected compounds in studies. Compound-specific physico-chemical and *in vitro* properties, such as lipophilicity, plasma protein binding, molecular mass, and plasma clearance have been applied and time-concentration profiles have been simulated. In an initial exploration, PBPK simulations were tested with different sources of lipophilicity (measured logD, calculated logD, and calculated logP) and in vitro clearance (rat liver microsomal assay and hepatocyte assay) (see Table S1). For missing in vitro data, the respective values were predicted based on already available ML models.

**Table S1** Overview of the Data set and compound characteristics

| Property | Reported for #compounds | Minimum | Median | Maximum |
|---|---|---|---|---|
| | | **All compounds [8040]** | | |
| Molecular mass (Da) | 8040 compounds | 130.0 | 450.0 | 6870.0 |
| Hepatocyte assay clearance (mL/min/kg) | 3524 compounds | 0.24 | 31.85 | 67.20 |
| Microsomal assay clearance (mL/min/kg) | 2180 compounds | 15.07 | 32.31 | 68.94 |
| Plasma protein binding (%) | 2998 compounds | 5.66 | 92.30 | 99.96 |
| Lipophilicity, logP (calculated, log units) [a] | 7997 compounds | -11.05 | 2.95 | 11.33 |

| | | | | |
|---|---|---|---|---|
| Lipophilicity, logD (at pH 7.4, log units) | 744 compounds | -2.6 | 2.3 | 6.1 |
| Lipophilicity, logD (at pH 7.4, calculated, log units [b] | 7986 compounds | -9.0 | 2.5 | 9.0 |

[a] calculated using Biobyte[5], [b] calculated using MoKa[6]

Furthermore, available methods for calculating the partitioning coefficients (PK-Sim Standard, Rodgers and Rowland, Schmitt, Berezhkovskiy) and the cellular permeabilities (PK-Sim Standard and Charge dependent Schmitt) were examined. Though different calculation methods provided better predictions for different types of compounds, the combination of the Berezhkovskiy partitioning coefficient and the PK-Sim cellular permeabilities calculation methods yielded better predictions on average. This combination has been selected for the generic framework to predict the PK of compounds in the presented work.

The current work examines the applicability of ML for prediction of the input parameters for a PBPK model from various compound descriptors. While most of the physico-chemical required for the parametrization of a PBPK model can be calculated from compound structure, the available lipophilicity and the clearance values possess were not sufficient to provide a sufficiently high *a priori* prediction quality, and both input parameters have a strong impact on PBPK model performance. To train the ML on the parameter set that produces the optimal prediction results with the applied PBPK framework, lipophilicity and plasma clearance have been optimized by fitting the model to measured time-concentration profiles.

**Details on ML models used in Baseline-ML and Pure-ML**

Baseline-ML and Pure-ML used the following predicted ADME/PK data features: LogD at pH=2, LogD at pH=11, high-throughput solubility at pH=2.2, high-throughput solubility at pH=4.5, high-throughput solubility at pH=6.8, metabolic stability in rat liver microsome assay, metabolic stability in rat hepatocyte assay with 5% serum, metabolic stability in rat hepatocyte assay with 50% serum, binding to rat plasma protein, permeability in Caco2 cell line, *in vivo* clearance in rats (NCA, not used for Baseline-ML), volume of distribution at steady state in rat (NCA, not used for Baseline-ML). The data was predicted using separate multi-task Chemprop models trained with the same identical temporal splits. The training was done in the same manner as for Compartmental-ML and PBPK-ML (see below).

**Details on ML models used in Compartmental-ML and PBPK-ML**

The ML models for both Compartmental-ML and PBPK-ML were trained using default hyperparameters in the Chemprop package (version 1.5.2) in python.[7] Each model was an ensemble of five individual neural network instances. For early stopping, a scaffold-based scheme was used to split the training data (90/10). All models were trained for up to 30 epochs and the best instance on the validation split used for early stopping was used to predict the corresponding test set in a temporal splitting scheme. In addition to graph convolutions, the models used RDKit features as global descriptors which are available within the Chemprop package. For Compartmental-ML, compounds were fitted either as one- or two-compartment. Data from the following experiments were used as auxiliary task in the multi-task model for both Compartmental-ML and PBPK-ML: LogD at pH=2, LogD at pH=11, high-throughput solubility at pH=2.2, high-throughput solubility at pH=4.5, high-throughput solubility at pH=6.8, metabolic stability in rat liver microsome assay, metabolic stability in rat hepatocyte assay with 5% serum, metabolic stability in rat hepatocyte assay with 50% serum, permeability in Caco2 cell line, *in vivo* clearance in rats (NCA), volume of distribution at steady state in rat (NCA). Binding to rat plasma protein was used as auxiliary task in Compartmental-ML (while being a required input to the PBPK model in PBPK-ML).

Each training set contained only auxiliary data for compounds registered prior to the cut-off date for the respective temporal split. Hence, chemical structures were the only information the model used about test compounds to make predictions.

To predict the required input parameters to the PBPK model (lipophilicity and CL), we used a comparable approach as for Compartmental-ML. Instead of predicting the four Compartmental-ML parameters, the two input parameters for the PBPK models were predicted based on chemical structures, along with the auxiliary tasks in a multi-task ML model.

### Additional figures

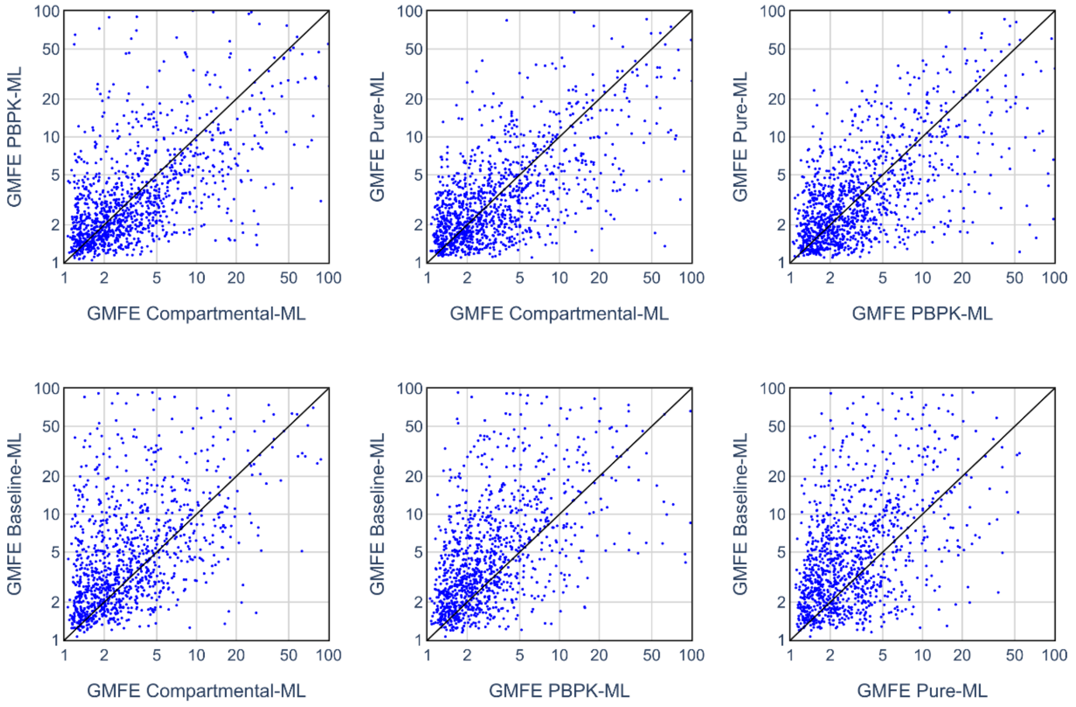


**Figure S1**. Each scatter plot shows the correlation between GMFE values obtained for two different PK profile prediction methods (within GMFE range [1,100]).

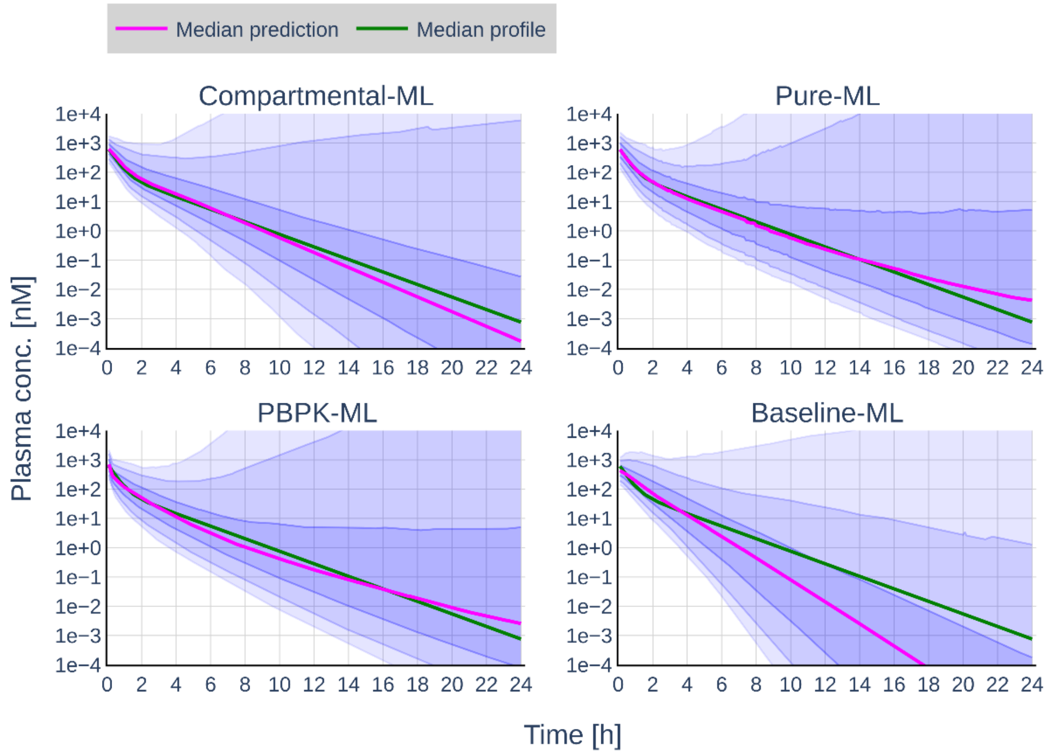


**Figure S2**. Visualization of prediction accuracy and bias over time. At each time point (0 to 24 hours in 6-minute intervals) the ratios of predicted concentrations to fitted concentrations were calculated. The plots show a typical profile generated from median parameters across the dataset (green line), the median deviation from a typical profile (50th percentile, pink line), as well as (from top to bottom) the 95th, the 90th, the 75th, the 25th, the 10th, the 5th percentile of fold errors for each method at the different time points.

### Additional tables

**Table S2.** Summary of GMFE values per method.

| | Compartmental-ML | PBPK-ML | Pure-ML | Baseline-ML |
|---|---|---|---|---|
| Median GMFE | 2.93 | **2.80** | 2.86 | 4.49 |
| % GMFE < 2 | **32.6** | 32.1 | 32.1 | 19.0 |
| % GMFE < 3 | 51.4 | **52.3** | **52.3** | 36.3 |
| % GMFE < 5 | 69.5 | 70.2 | **71.0** | 52.9 |
| % GMFE < 10 | 81.7 | 82.0 | **84.1** | 70.2 |
| % GMFE < 100 | 95.1 | 96.5 | **98.7** | 88.7 |
